# FH loss in RPE cells causes retinal degeneration in a human RPE-porcine retinal explant co-culture model

**DOI:** 10.1101/2021.07.26.453778

**Authors:** Angela Armento, Aparna Murali, Julia Marzi, Blanca Arrango-Gonzalez, Ellen Kilger, Simon J Clark, Katja Schenke-Layland, Charmaine A Ramlogan-Steel, Jason C Steel, Marius Ueffing

## Abstract

Age-related Macular degeneration (AMD) is a degenerative disease of the macula affecting the elderly population. Treatment options are limited, partly due to the lack of understanding of AMD pathology and the sparse availability of research models, that replicate the complexity of the human macula and the intricate interplay of the genetic, aging and life-style risk factors contributing to AMD. One of the main genetic risks associated with AMD is located on Complement Factor H (*CFH*) gene, leading to an amino acid substitution in the FH protein (Y402H). However, the mechanism of how this FH variant promotes the onset of AMD remains unclear. Previously, we have shown that FH deprivation in RPE cells, *via CFH* silencing, leads to increased inflammation, metabolic impairment and vulnerability towards oxidative stress. In this study, we established a novel co-culture model comprised of *CFH* silenced RPE cells and porcine retinal explants derived from the visual streak of the porcine eyes, closely resembling the human macula. We show that retinae exposed to FH-deprived RPE cells show signs of retinal degeneration, with rod cells being the first cells to undergo degeneration. Moreover, *via* Raman analyses, we observe that the main changes involve the mitochondria and lipid composition of the co-cultured retinae upon FH loss. Interestingly, the detrimental effects of FH loss in RPE cells on the neuroretina were independent of glial cell activation and external complement sources. Moreover, we show that the co-culture model is also suitable for human retinal explants, and we observed a similar trend when RPE cells deprived of FH were co-cultured with human retinal explants from a single donor eye. Our findings highlight the importance of RPE derived FH for retinal homeostasis and provide a valuable model for AMD research.

## 1. Introduction

Age related macular degeneration (AMD) is a complex multifactorial disease that severely compromises the visual acuity and eventually leads to irreversible blindness if left untreated [1, 2]. It is the leading cause of blindness among elderly population globally, and the numbers are expected to increase over the next few years due to the increasing aged population [3]. The region of the eye affected in AMD is the macula, the central region of the retina with the highest number of photoreceptors (PR), which are located on top of RPE cells that themselves sit on top of a thin membrane called Bruch’s membrane (BM), which separates the neuroretina from the choroid, a network of blood capillaries [4]. Up to now, no satisfactory treatment has been discovered for the more common late-stage form of AMD, which is characterized by RPE geographic atrophy (GA). The reasons for that can be associated with the complexity of AMD pathology, which involves a variety of risk factors (ageing, genetic predisposition, life style) [5] and the complexity of a multi-layer/multi-cellular structure like the retina. Moreover, the research models used so far, do not fully replicate important aspect of the macula [6]. For example, mono-cellular *in vitro* studies do not take into consideration the interactions between cell types, and animal models, though largely used for retinal studies, do not replicate the specific properties of the macula, which is unique to humans [7]. A lack of macula-like research models makes it very difficult to investigate the order of events in AMD pathology. Moreover, it is unclear whether rods or cones PR die first and, most importantly, the mechanism behind the cell death of either one of them. Studies from Curcio *et al*. proposed that rod cell death may precede cone cell death in AMD [8], since in the para-foveal region during the early stages of AMD, the rod PR undergo extensive apoptosis compared to the cones.

Although AMD is triggered by multiple genetic and environmental factors, a large portion of the genetic risk falls into genes of the alternative pathway of the complement system. In particular, one of the most strongly associated genetic variant with AMD risk (rs1061170) resides in the *CFH* gene, which encodes the factor H and factor H-like proteins: potent regulators of complement activation [9, 10]. This genetic variant leads to a coding change in the FH/FHL-1 proteins where a tyrosine residue is replaced by a histidine residue at position 402 (residue 384 in the mature protein) and is referred to as the Y402H polymorphism. Furthermore, secreted/extracellular FH/FHL-1 has been described to possess additional functions and the Y402H polymorphism has been related to different aspects of AMD pathology [4]. For instance, RPE cells are constantly exposed to oxidative stress due to the fact that the retina consists of an highly oxidized environment and RPE cells are responsible for the degradation of oxidized phospholipids shed by PR [11]. In this context, FH plays an anti-oxidant protective role [12] directly binding oxidized lipids [13]. The 402H high-risk variant of FH protein loses this protective function, probably due to the decreased affinity for oxidized lipids [13], resulting in a reduced ability of the RPE cells to neutralize oxidized lipids, which then accumulate and contribute to drusen formation and thereby promote oxidative stress and inflammation [14, 15]. Moreover, RPE cells exist in a complex co-dependent metabolic relationship with PR, since the PR rely on RPE cells for the supply of nutrients and a bioenergetic crisis of RPE cells has been described to be part of AMD pathology [16].

Recently, a novel function for intracellular FH has been described in several cell types and place intracellular FH as an important regulator of cell homeostasis [17, 18]. iPSC-RPE cells, carrying 402H high-risk variant of FH, show enlarged mitochondria[19] and reduced mitochondria activity [20], but whether the effects depends on intracellular FH or extra-cellular FH was not investigated. Our previous studies show that the loss of endogenous/intracellular FH in RPE cells leads to a phenotype very similar to iPSC-RPE cells carrying 402H high-risk variant of FH. In detail, hTERT-RPE1 cells deprived of endogenous FH *via CFH* silencing, are more vulnerable to oxidative stress, show reduced metabolic capacity [17] and modify the microenvironment toward an inflammatory state [21], all predisposing features for AMD.

This study investigates the impact that endogenous loss of FH in RPE cells has on the retina in a novel hybrid co-culture model composed of human RPE cells and porcine/human retinal explants. The presence of RPE cells helps to maintain the retinal architecture *ex-vivo*, as shown by Mohlin *et al*. when co-culturing ARPE19 cells and porcine retinal explants [22]. Furthermore, porcine retina has morphological and functional similarities to the human retina, including its cone-rich visual streak, an avascular retina area rich in cone PR, similar to the human macula being an attractive model to study retinal function and disease [23]. Porcine retinal explants have been successfully used to study other retinal disease mechanisms, such as oxidative stress- or hypoxia-mediated damage, characteristics of several retinal degenerative disease, including AMD [24, 25].

In this study, we employed retinal explants obtained exclusively from the visual streak region of the porcine retina, to mimic as much as possible the human macula. We show that retinae exposed to RPE cells deprived of FH undergo a faster photoreceptor cells loss and it appears that the rod cells are the first cells to undergo degeneration. Moreover, *via* Raman analyses, we observe that the main changes in the retina appear to be in the mitochondria and lipid composition of the co-cultured retinae upon FH loss. Furthermore, we prove that the co-culture model is also suitable for human retinal explants, and we observed a similar trend when RPE cells deprived of FH were cultured from human retinal explants from a single donor eye. Our findings highlight the importance of RPE-derived FH for retinal homeostasis and provide a valuable model for AMD research.

## 2. Materials and Methods

### Cell culture

Human retinal pigment epithelium (RPE) cell lines hTERT-RPE1 cells were obtained from the American Type Culture Collection (ATCC). The cells were cultured in medium containing 1:1 DMEM F12: Hams F12 (Gibco, Germany) with additional supplementation of 10% fetal calf serum (FCS; Gibco, Germany), penicillin (100 U/ml) and streptomycin (100µg/ml). These RPE cells were seeded at the rate of 100,000 cells/transwell in a 12mm polyester transwell with 0.4micron pore size and were allowed to grow overnight at 37°C incubated with 5% carbon dioxide. The *CFH* gene in the RPE cells was silenced by siRNA-mediated transfection according to the manufacturer’s instructions using Viromer Blue reagent (Lipocalyx, Germany). The siRNA (IDT technologies, Belgium) mixture was prepared using 3 different silencing RNAs specific for *CFH* (siCFH) and one siRNA for negative control (siNeg) in the Viromer Blue buffer. After 24 hours of silencing, the media was changed, and the cells were used for co-culture.

### Porcine retinal explants

The porcine eyes were obtained from the local slaughterhouse. All the experiments conducted with porcine retinal explants were approved by the local institutional ethics committee. The explants were prepared from 6-month-old pigs weighing on an average 100kg within 2-3 hours of enucleation as previousely described [26]. They were euthanized by electrocution and the enucleated eyes were transported in ice directly to the research facility. The eyes were immersed in 70% ethanol, following which the muscle tissue was removed using sterile Westcott scissors in a laminar air flow chamber. A small incision was made using a sterile blade, and the anterior segment of the eye was removed including the cornea, lens, iris and ciliary body. Sharp forceps were used to break the vitreous free from the eye cup and the vitreous was also removed. In the avascular zone, two cone-rich zones in the retina were identified and dissected based on the pattern of blood vessels (as shown in Figure S1). The retina was then carefully peeled off from the RPE-choroid-sclera layers and placed on the transwell containing RPE cells, with the photoreceptor side facing down towards the RPE cells. From each porcine eyecup, 2 cone rich retinal explants were obtained to have an internal control for each pig: one explant was exposed to control siNeg RPE cells and one explant to siCFH RPE cells. The RPE-retina was co-cultured in medium containing 1:1 DMEMF12: Hams F12 with B27 supplement (Life Technologies), penicillin (100 U/ml) and streptomycin (100µg/ml) for 72 hours.

### Human retinal explants

The human donor eye was obtained from the human tissue biobank of the Eye Clinic Tübingen (ethic 324/2013BO1). Prior informed consent was obtained from the donor and the anonymity of the donor was maintained. The human donor was diagnosed with uveal melanoma and the enucleation surgery was performed at the University Hospital Tübingen in Tübingen, Germany. The human eyecup was transported to the laboratory in ice within 30 minutes of enucleation and experiments were performed accordingly to ethic 185/2021BO2. Using sterile forceps, the vitreous was removed. The retina was randomly dissected to obtain 6 explants, excluding the region surrounding the dissected tumor. The human retinal explants were co-cultured with negative control (siNeg) and *CFH* silenced (siCFH) hTERT-RPE1 cells grown on transwell as described above. The hTERT-RPE1 cells-human retina explants were co-cultured *ex-vivo* in in medium containing 1:1 DMEMF12: Hams F12 with B27 supplement (Life Technologies), penicillin (100 U/ml) and streptomycin (100µg/ml) for 72 hours.

### Immunohistochemistry

The co-cultured RPE-retina were washed with 1X PBS for 5 minutes and fixed using 4% PFA for 45 minutes in room temperature. After that a sucrose gradient was performed by incubating the samples in 10%, 20%, 30% sucrose to ensure cryopreservation of the co-cultured tissue. The tissues were then snap-frozen in liquid nitrogen using Tissue-Tek Optimal cutting temperature compound. The cryo moulds are stored at -20°C until sectioning. Using a cryostat, 12 to 14-micron thick retina-RPE co-culture sections were obtained. The cells were then permeabilized and blocked for 1 hour with 0.1% Triton X-100,10% normal goat serum and 1% bovine serum albumin in PBS. The sections were then stained overnight at 4°C in the respective primary antibodies (anti-Rhodopsin, MAB5316 Sigma-Aldrich; anti-M-Opsin, AB5405 Sigma-Aldrich; anti-S-Opsin, AB5407 Sigma-Aldrich; anti-Iba-1, 019-19741 Wako Chemicals; anti-GFAP, G-3893 Sigma-Aldrich; anti-Bestrophin-1, AB2182 Sigma-Aldrich). After overnight incubation, the sections were washed thrice with PBS to remove excess primary antibody. The recommended concentration of secondary antibodies (Goat Anti-Rabbit Alexa Fluor 488, A-11034 and Goat Anti-Mouse Alexa Fluor 568, A-11031, Invitrogen) were added and the sections were incubated for 1 hour. The sections are again washed thrice in PBS and stained with DAPI for 5 minutes and mounted with Fluoromount-G solution (17984-25, Electron Microscopy Sciences, Pennsylvania, USA). Using a Zeiss Axio Imager Z1 ApoTome Microscope, Z-stack images were obtained and the images were analyzed using Zen Blue image analysis software.

### Image analysis

For each sample, 3 sections were selected and for each section, 3 different retinal regions were imaged to obtain 9 images for each sample. To calculate the overall retinal thickness, thickness of ONL and number of rows in ONL, 12 data points were obtained for each image to get a uniform coverage of an entire image. All the images were taken with uniform exposure time and the fluorescence of all the images are also brought to uniform values within the Zen Blue software, so as to obtain a comparable mean fluorescence intensity value. To obtain the density of M- and S-cones, 3 * 100 micron regions were selected in each image and the average of 9*3 = 27 data points was considered as the number of cones in 100µm. As the majority of the cones are positioned on the outermost layer of the ONL in healthy conditions, displaced cones were counted as cone positive nuclei found 3 rows above the outer most layer of the ONL.

### Raman microspectroscopy

Cryosections obtained from sample preprocessing for immunohistochemistry were thawed, rinsed with PBS and kept hydrated during the whole measurement. Raman images were recorded using a WiTec 300R alpha Raman microspectroscope (WiTec GmbH, Ulm, Germany), equipped with a green laser (532 nm), a 63x dipping objective (Carl Zeiss AG, Jena, Germany) and a spectrograph with a 600 g/mm grating and a CCD camera. Spectral maps were acquired as 100×300 µm cross-sections with a pixel resolution of 1×1 µm, a laser power of 50 mW and an integration time of 0.1 s/spectrum. In addition, a reference spectrum of docosahexaenoic acid (DHA, #53171, Sigma Aldrich) was generated as an average spectrum from spectral maps of the pure substance (1×1 µm spatial resolution, 0.1 s integration time, 50 mW laser power).

### Raman data analysis

Before further analysis, the acquired spectral maps were preprocessed in the WiTec *Project Five* Software (5.4, WiTec GmbH). Data underwent cosmic ray removal, were baseline corrected (shape algorithm) and normalized (area to 1). True component analysis (TCA) was applied to generate false color-coded intensity distribution heatmaps of major tissue structures as previously described [27]. For quantitative analysis, intensity heatmaps were extracted for each component and mean GVIs were calculated in ImageJ and normalized to the cell number. Analyses were focussed on the ONL region. A number of 3 spectral maps per sample was analyzed.

Furthermore, single spectra (n=100/image) were extracted from the lipid-assigned regions for in-depth analysis of the molecular composition via principal component analysis (PCA) performed in The Unscambler X (Camo Analytics AS, Oslo, Norway). The application of PCA for the analysis of spectral data has been described elsewhere [28]. In brief, principal component (PC) score values were compared between the two groups to identify differences in their molecular composition. The underlaying molecular information can be retrieved from the corresponding PC loadings plot, indicating relevant spectral features as positive or negative peaks.

### RNA isolation, cDNA synthesis and Qrtpcr

RPE cells pellets were collected and RNA was extracted using Tri-Fast reagent according to the manufacturer instructions. The concentration of the isolated RNA samples was assessed using Nanodrop spectrophotometer. cDNA synthesis was performed *via* reverse-transcription of 2µg of RNA using M-MLV Reverse Transcriptase (Promega (Wisconsin, USA). cDNA was used to analyze silencing efficiency by qRT-PCR employing iTaq Universal SYBR Green Supermix (Bio-Rad Laboratories, USA) along with gene-specific forward and reverse primers (10 μM, PRPL0 fw 5′-GGA GAA ACTGCT GCC TCA TATC -3, PRPL0 rev 5′-CAG CAG CTGGCA CCT TAT T -3′, *CFH* fw 5′-CTG ATC GCA AGAAAG ACC AGT A -3′, *CFH* rev 5′-TGG TAG CACTGA ACG GAA TTAG -3′).

### C3b ELISA

Based on manufacturer’s instructions, the concentration of C3/C3b in culture supernatants were analysed by ELISA assay (Abcam, UK). Samples were loaded undiluted along with standards and controls in 96 well-plates coated with specific C3b antibody and the assay was performed according to the manifacturer instructions. Absorbance was immediately measured at 450nm using a Spark multimode microplate reader (Tecan, Switzerland). In order to correct optical imperfections, subtraction readings were taken at 570 nm.

### Statistical analysis

The graphs were plotted as mean ± SEM using Graph Pad Prism version 8 and the statistical analysis was performed using SPSS version 2.2 and Graph Pad Prism version 8. Quantification of western Blot signals and fluorescent images was done using ImageJ and Zen Blue Image analysis software respectively. Unpaired and paired t-test were used to assess the differences in the means between different groups. A P-value less than 0.05 is considered statistically significant.

## 3. Results

### 3.1. Porcine retinal explants exposed to FH-deprived RPE cells shows signs of degeneration

Our previous work showed that the homeostasis of RPE cells deprived of FH is strongly impaired. This study aims to evaluate the impact that RPE cells deprived of FH have on the neuroretina. We used a co-culture system comprising human hTERT-RPE1 cells and porcine retinal explants obtained from the porcine visual streak cone-rich area to study the impact of those impaired RPE cells on the retina. Thus, hTERT-RPE1 cells were either silenced for the *CFH* gene (siCFH) or with negative control (siNeg), as already shown in Armento *et al*. and Figure S2. As shown in Fig. 1A, retinal explants were collected and placed in direct contact with the RPE cells in the transwell with the PR side facing the RPE cells, which mimics the in-vivo condition (shown in Fig. 1A). The porcine retinal explants co-cultured for 3 days *ex-vivo* with FH-deprived hTERT-RPE1 cells (siCFH) exhibited signs of degeneration compared to retinal explants co-cultured with siNeg hTERT-RPE1 cells (Fig. 1B-C). In detail, overall retinal thickness was reduced (176.8 ± 5.3 µm vs 146.4 ± 8.2 µm, Fig. 1D), the thickness of the outer nuclear layer (ONL) was smaller in siCFH co-cultured retinae (51.7 ± 2.7 µm vs 38.9 ± 1.2 µm, Fig. 1E) and the number of cell rows in the ONL was lower (8.4 ± 0.5 vs 6.1 ± 0.2; Fig. 1F). In order to ensure that the effects observed are dependent on the status of the RPE cells and not dependent on the loss of RPE cells, which could occur upon FH loss, we exposed the retinal explants to different concentration of hTERT-RPE1 cells (Figure S3). Our results indicate that the porcine retina co-cultured with 50,000 hTERT-RPE1 cells (seeding density) had a similar retinal structure to that of the porcine retina co-cultured with 100,000 hTERT-RPE1 cells (seeding density also employed for siNeg-siCFH experiments). Interestingly, upon seeding 200,000 hTERT-RPE1 cells, it had a negative impact on the retinal parameters investigated and in particular, ONL thickness was significantly reduced (Figure S3). From this data, it is evident that even in case of reduced number of cells in siCFH hTERT-RPE1 condition; this could not affect the co-cultured porcine retinal architecture.

**Figure 1:**
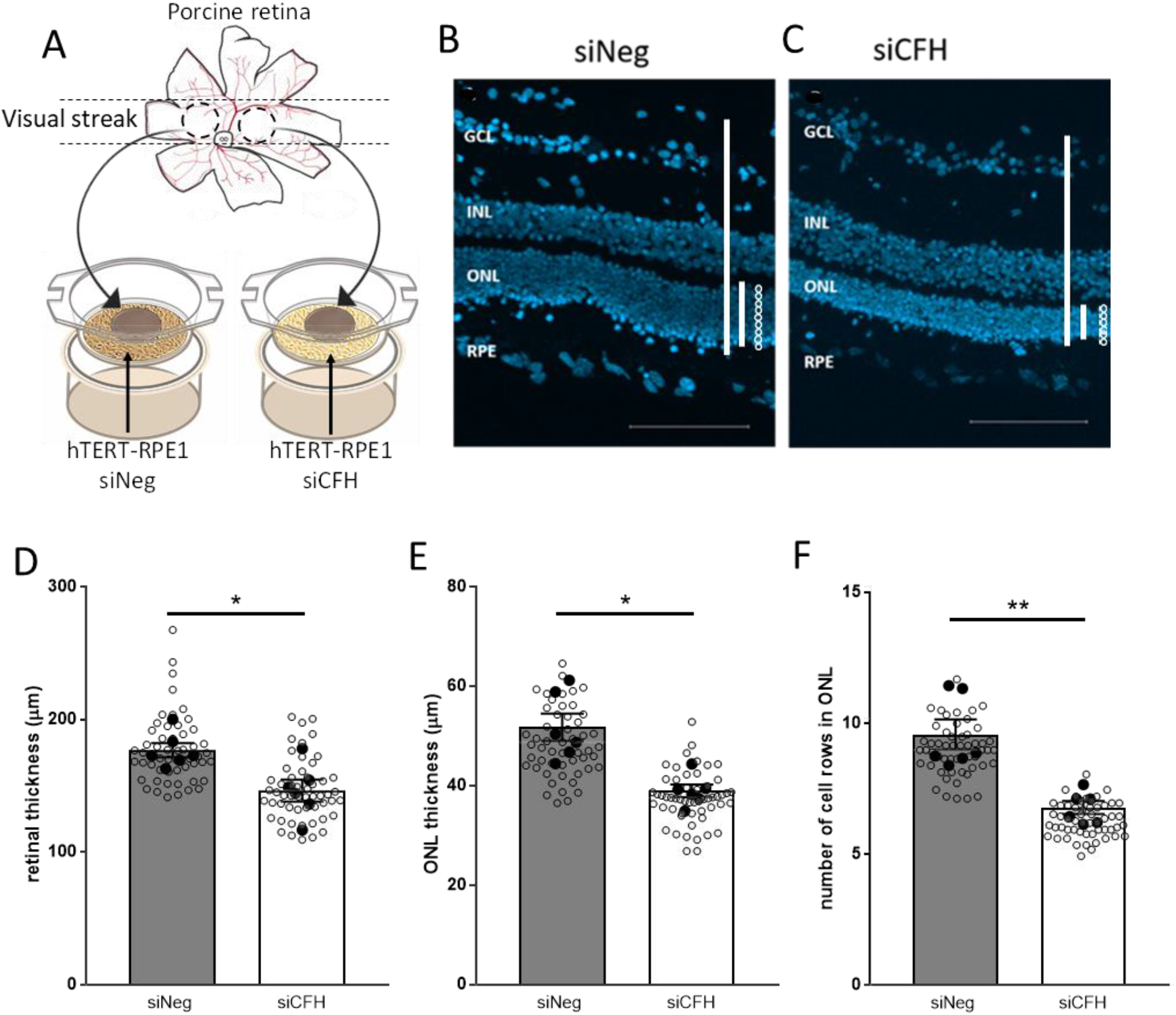
FH loss-mediated changes in the co-cultured porcine retina. **A** Schematic representation of the co-culture system, in which two retinal explants from the visual streak of porcine donor eyes are exposed to either siNeg or siCFH hTERT-RPE1 cells for 72 hours. **B-C** The porcine retinal architecture was assessed with the help of DAPI, a nuclear staining dye. Representative images of siNeg coculture (B) and siCFH co-culture (C) are shown. Lines and circles highlight the parameters extrapolated from these images and quantified in **D-F**: the overall thickness of the porcine retina (D), the thickness of the ONL (E) and the number of rows in the ONL (F). Abbreviations: GCL – Ganglion cell layer, INL – Inner nuclear layer, ONL – Outer nuclear layer, RPE – Retinal pigmented epithelium cells, siNeg – silencing negative control, si*CFH* – silencing *CFH*. Mean ± SEM is shown. N= 6 biological replicates. *p ≤ 0.05; **p ≤0.005. Scale bar - 100µm.

### 3.2. FH loss in RPE cells mediates rod cell degeneration in the co-cultured porcine retina

As the second step, we focused on identifying which cell type was mainly affected by FH loss in RPE cells. The porcine rod and cone PR were immune-labeled with rhodopsin and opsin antibodies respectively after 3 days in co-culture with either siNeg or siCFH hTERT-RPE1 cells (Fig. 2A-B). Cell counting revealed a significant reduction in the total number of cells in the ONL, indicating a loss of PR cells (253.3 ± 14.5 cells vs 200 ± 6.6 cells, Fig. 2C). In order to understand whether rod or cone PR cells are degenerating first, we analyzed the specific rhodopsin and opsin staining. We did not observe any reduction in the number of total M-cones (Fig. 2 D), indicating that a loss of rods causes the reduced number of cells in ONL. Although the number of M-opsin positive cells (Fig. 2D) were similar in both groups, the distributions of M-cones were disrupted in the absence of FH in the RPE cells and more displaced cones were counted when the retinae were co-cultured with siCFH hTERT-RPE1 cells (Fig. 2E).

**Figure 2:**
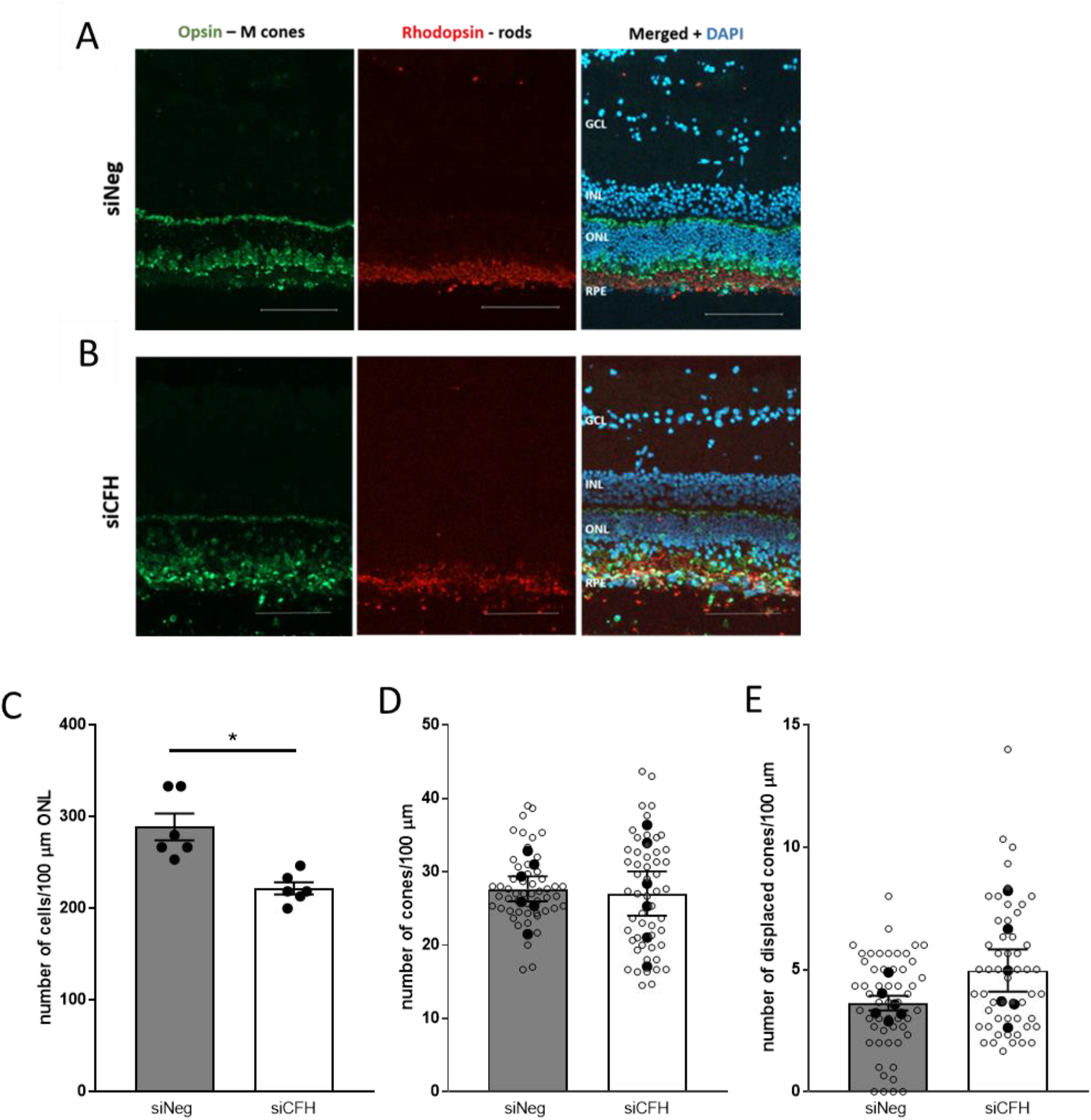
Effect of FH loss in RPE cells on porcine PR in co-culture. **A-B** Retinal explants from the visual streak of porcine donor eyes are exposed to either siNeg (A) or siCFH (B) hTERT-RPE1 cells for 72 hours. Explants cryosections were stained for Opsin (green) and Rhodopsin (red) to differentiate between cones and rods and DAPI was used as counterstaining for nuclei. **C-E** Quantification of retinal parameters extrapolated from A-B: number of cells in ONL (C), number of cones in ONL (D) and number of displaced coned (E). Abbreviations: GCL – Ganglion cell layer, INL – Inner nuclear layer, ONL – Outer nuclear layer, RPE – Retinal pigmented epithelium cells, siNeg – silencing negative control, si*CFH* – silencing *CFH*. Mean ± SEM is shown. N= 6 biological replicates. *p ≤ 0.05; **p ≤0.005. Scale bar - 100µm.

Our previous study shows that FH deprivation in RPE cells leads to elevated secreted levels of C3. To assess whether C3 directly mediates retinal damage, we supplemented the culture medium with purified C3. Exogenous addition of purified C3 to the porcine retina co-cultured with either siCFH or siNEG hTERT-RPE1 cells had no impact on the retinae and the observed changes in the absence of C3 were maintained (Figure S4). Moreover, we evaluated whether the addition of purified FH could overcome the absence of endogenous FH in RPE cells. Exogenous addition of purified FH protein to the co-culture media did not induce any effect in the porcine retina compared to samples without additional FH (Figure S4). This data indicates that the C3 and FH proteins purified from the human serum do not directly influence on the porcine retina.

Moreover, RPE cells deprived of FH show an increase in inflammatory cytokines production [21] and since inflammation is a sign of degeneration [29], we assessed the levels of inflammatory cells in the retinal explants. After 3 days of co-culture, we evaluated the number of Iba1 positive cells as an indicator of microglial cells in retinal explants co-cultured with both siNeg and siCFH hTERT-RPE1 cells (Figure S5) and we did not observe any difference in any of the retinal layers (Figure S5). In addition, the retina presents a specialized type of microglia cells, the Muller cells, whose activation in the porcine retina was marked by anti- glial fibrillary acidic protein (GFAP) antibody (Figure S5). However, we did not find any significant differences in the mean fluorescence intensity of GFAP in the retina co-cultured with either siNeg or siCFH hTERT-RPE1 cells (Figure S5).

### 3.3. RPE cells deprived of FH have a negative impact on the mitochondria in the cells of the ONL

In order to understand the type of damage that RPE cells deprived of FH provoke in the retina, we performed Raman imaging for marker-independent and molecular-sensitive analyses. Raman microspectroscopy allows to generate a specific molecular fingerprint based on the assignment of spectral bands to distinct molecular groups. In this way, we were able to identify and localize different subcellular structures in the samples (Fig. 3A). TCA allowed to distinguish nuclei, mitochondria and two different groups of lipids according to their specific spectral signatures (Fig. 3B). Nuclei showed specific DNA-related bands at 792, 1098 and 1580 cm-1, mitochondria were characterized by their cytochrome C signals at 750, 1130 and 1589 cm-1 and the lipid components showed peaks at 720, 1450, 1655 and 2850 cm-1 correlating to phospholipids and fatty acids. First, the mitochondria signal was analyzed in detail. Since the ONL is the layer of the neuroretina affected in AMD, we specifically quantified cytochrome c signal intensities within this region (Fig 3C) and demonstrated a loss in cytochrome c signal in the ONL of the co-culture samples with the siCFH RPE cells (Fig. 3D). The identified cytochrome c spectrum (Fig. 3E) corresponds to the signature of the reduced, bound form found in healthy cells and has a different Raman signature compared to oxidized cytochrome c [30, 31], which is released into the cytosol upon apoptosis. The reduction of cytochrome c in the mitochondria in the ONL of the FH deprived group indicates lower mitochondrial activity and might be a consequence of oxidative stress.

**Figure 3.**
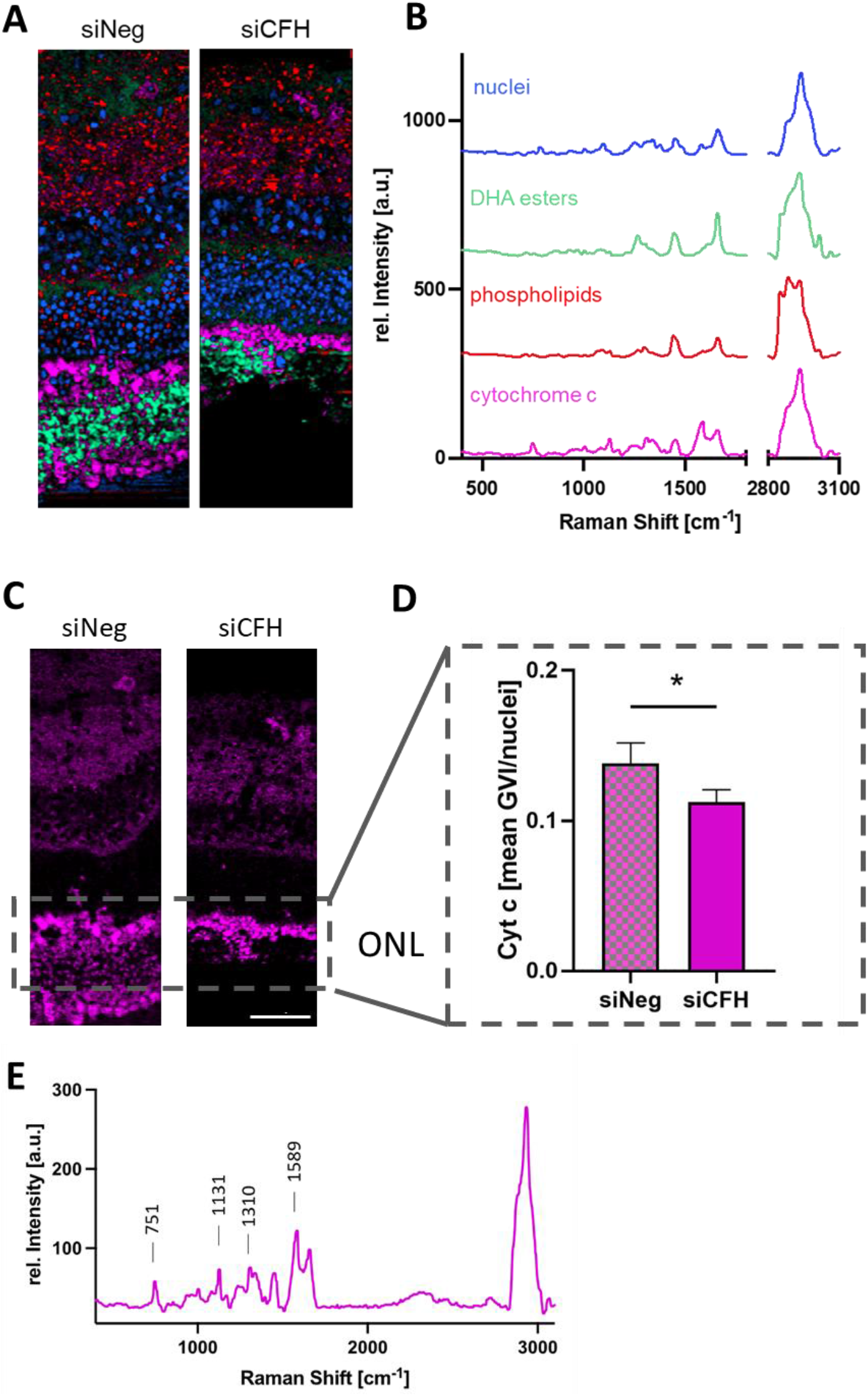
**A, B** True component analysis (TCA) was performed on the spectral maps of siNeg and siCFH samples to generate intensity distribution heatmaps (A). According to different spectral signatures (B) pixels assigned to nuclei (blue), cytochrome c (pink) and two different lipid components (red and light green) could be identified and localized. **C, D** Quantitative analysis of the mitochondrial activity of the ONL region based on the cytochrome c intensity distribution images (C) and calculation of the mean GVI/cell (D). **E** Raman signature of the mitochondrial activity TCA component. ONL – Outer nuclear layer, siNeg – silencing negative control, siCFH – silencing CFH. Data are presented as mean ± SD. N= 3 biological replicates. *p ≤ 0.05; Scale bar – 50 µm.

### 3.4 Retinae cultured with siCFH RPE cells showed signs of lipid oxidation in the ONL

Raman image analysis via TCA identified two lipid components that showed different distribution patterns within the retina (Fig. 3A). Whereas the first component (green) was predominantly localized in the ONL, the second lipid signal (red) was rather associated with the inner retina. The ONL assigned lipids were quantified based on their signal intensities (Fig. 4A, B) and showed no significant difference between both groups. To identify the lipid subtype, average spectra from siCFH and siNeg were compared to a reference spectrum of docosahexaenoic acid (DHA), which is one of the most abundant phospholipid in the outer ONL [32]. The spectral overlay (Fig. 4C) demonstrated similar spectral signatures between DHA and the ONL lipids. For in-depth analysis and in addition to quantitative analysis, qualitative analysis via PCA was performed to investigate if changes in lipid composition are induced upon FH deprivation. Comparison of the single spectra in the PCA scores plot (Fig. 4D) and analysis of the average PC score values (Fig. 4E) demonstrated a significant shift in the spectral signatures of both groups described by PC-5. The assigned spectral changes were exploited as negative (siNeg) or positive (siCFH) peaks in the corresponding PC-5 loadings plot (Fig. 4F). In comparison to siCFH samples, the control samples had more intense bands at 1270 and 1655 cm-1 and a higher 2850 cm-1 signal, whereas the siCFH group was characterized by an increase of the 2940 cm-1 band. The identified spectral features correspond to C-H and C=C bonds in lipid molecules[33] and have been reported before in the context of lipid oxidation, where oxidative processes have been linked to decreases in the 1270 cm-1 and 2840 cm-1 region and to an increase of signal intensity at the 2940 cm-1 band [34, 35]. These findings correlate to our observations and indicate the relevance of lipid oxidation in DHA esters as one of the destructive processes induced by siCFH RPE cells. Furthermore, we analyzed the Raman images of the second lipid component, which was mainly localized in the inner retina and contained phospholipid-assigned spectral features (Fig. S6B). There was no quantitative difference detected in the lipid signal intensities of both groups (Fig. S6 A, C) and no significant structural alteration occurred, as demonstrated by no separation in any of the calculated PCs (Fig. S6 D).

**Figure 4:**
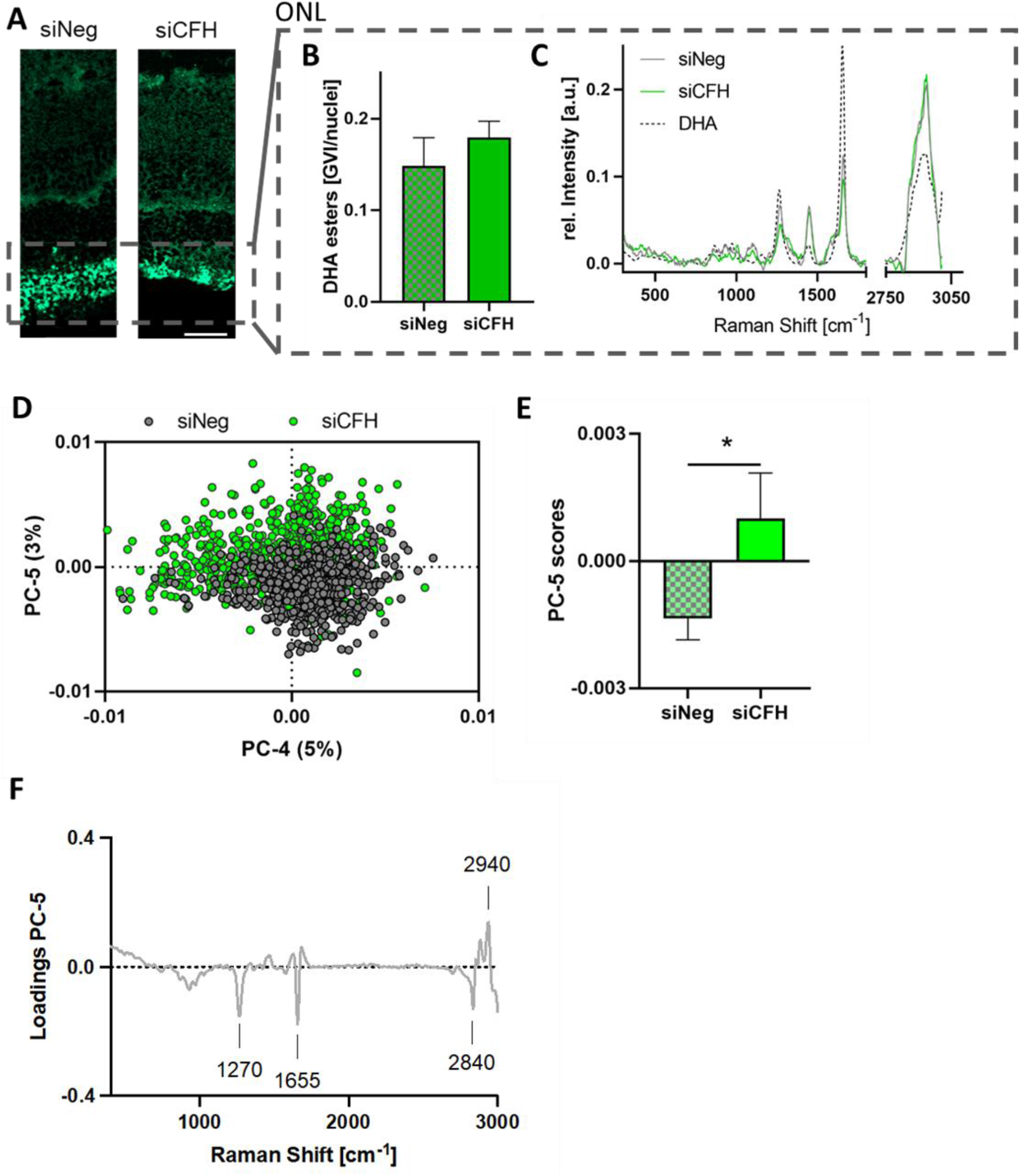
**A** TCA intensity distribution heatmaps of the first lipid feature showed a dominant localization in the ONL region. **B** Quantification of lipid signal/cell in the ONL. **C** Overlay of a DHA reference spectrum (black, dashed) with the average siCFH (green) and siNeg (grey) spectra from the lipids in the ONL region. **D-F** Principal component analysis (PCA) of single spectra (n=100 per sample) extracted from siNeg (grey) and siCFH (green) ONL regions demonstrates a clustering in the PCA scores plot (D) with a significant difference described by PC-5 (E). The most relevant spectral signatures are demonstrated in the loadings plot (F). Data are represented as mean ± SD, n=3, *p ≤ 0.05, scale bar – 50 µm.

These results lead to the conclusion that the FH deprived RPE cells do impact the lipids in the ONL – most likely by oxidative processes on the DHA esters - but do not induce further alterations on the phospholipids of the inner retina layers.

### 3.5 FH loss in RPE cells leads to degeneration in human retinal explants

Ultimately, we investigated in a human RPE-human retinal explant model whether the effects observed in porcine retinal explants are conserved in the human retina in response to FH-deprived RPE cells. Due to the scarce availability of human eye donors, explants from one donor were collected and exposed to either siNeg or siCFH hTERT-RPE1 cells for 72 hours (Fig. 5A-D) and the same retinal parameters were analyzed as for the porcine retinal explants. First of all, we confirmed that a co-culture system including human RPE cells and human retinal explants from different sources is possible. However, we did not observe any changes in retinal thickness (Fig. 5 E). However, ONL thickness of the retinal explants co-cultured with siCFH hTERT-RPE1 cells was slightly reduced compared to siNeg controls (21.9 ± 6.1. µm vs 20.6 ± 2.8 µm; Fig. 5 F). Similarly, the number of ONL cell rows in the retina co-cultured with siCFH hTERT-RPE1 cells was lower (4.6 ± 0.5 vs 2.6 ± 0.8, Fig. 5 G). Although these reductions in the number of rows in ONL and the ONL thickness are not statistically significant, this decline might be indicative of a loss of PR, most likely of rod PR since also in the case of human retinal explants, the number of cones was not affected at this time point (Fig. 5 H). Additionally, the hypothesis that rod degeneration precede cones degeneration in this model is supported by the observation that rod outer segments are strikingly smaller in retinae co-cultured with siCFH compared to siNeg hTERT-RPE1 cells (Fig. 5A-B).

**Figure 5:**
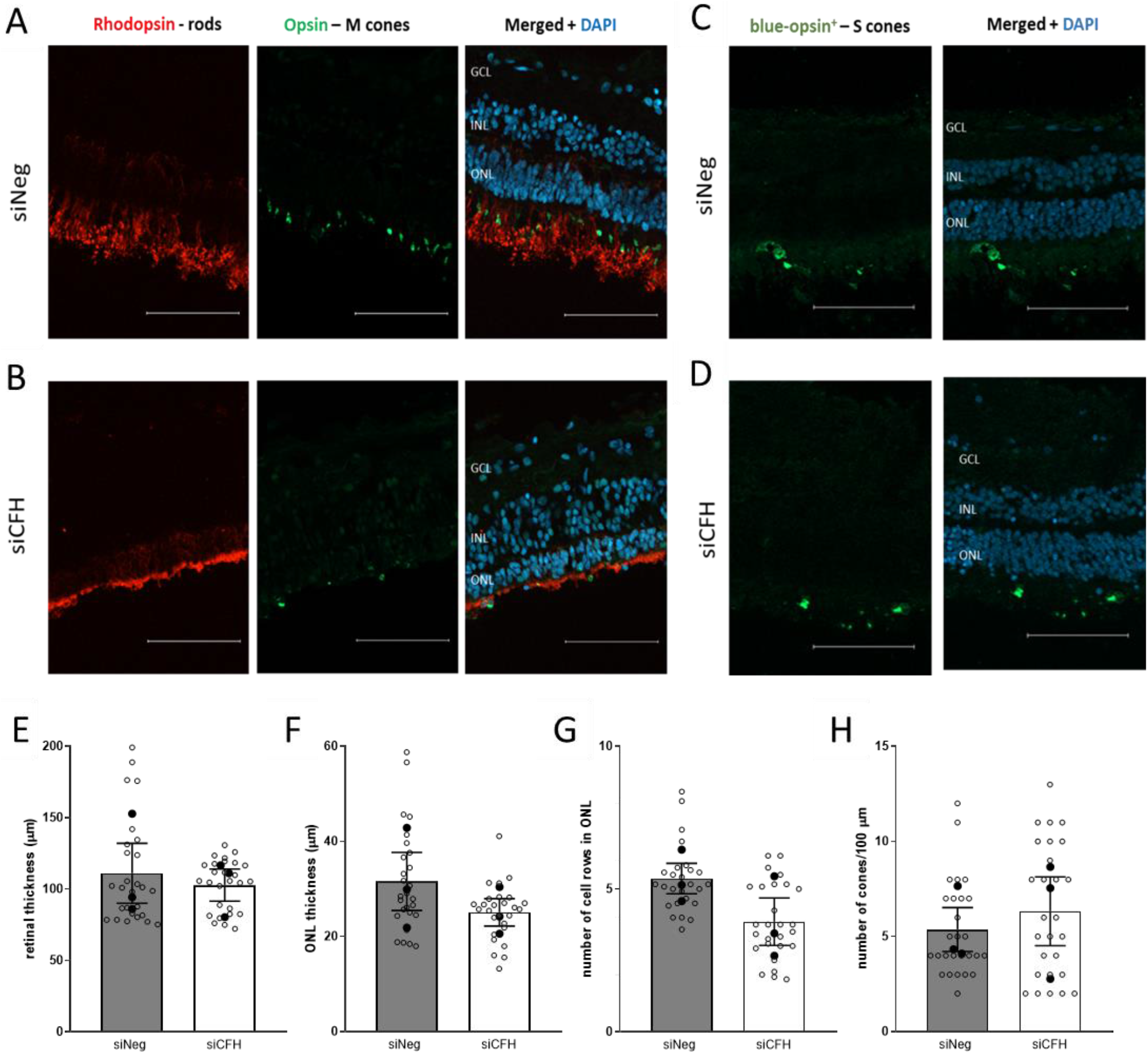
Changes mediated by FH loss in RPE cells in the co-cultured human retina. Human retinal explants from one single donor eye are exposed to either siNeg (A-C) or siCFH (B-D) hTERT-RPE1 cells for 72 hours. **A-B** Explants cryosections were stained for M-Opsin (green) and Rhodopsin (red) to differentiate between cones and rods and DAPI was used as counterstaining for nuclei. **C-D** Explants cryosections were stained for S-Opsin (green) and DAPI was used as counterstaining for nuclei to identify S-cones. **E-H** Quantification of retinal parameters extrapolated from A-D: retinal thickness (E), ONL thickness (F), number of cell rows in ONL (G) and number of cones in ONL (H). Abbreviations: GCL – Ganglion cell layer, INL – Inner nuclear layer, ONL – Outer nuclear layer, RPE – Retinal pigmented epithelium cells, siNeg – silencing negative control, si*CFH* – silencing *CFH*. Mean ± SEM is shown. N= 3 biological replicates from a single donor eye. *p ≤ 0.05; **p ≤0.005. Scale bar - 100µm.

## 4. Discussion

AMD is a complex multi-factorial disease, whose aetiology and pathology are not yet fully understood [36]. Reliable retinal experimental models are pre-requisites for exploring the pathological changes in this degenerative disease. In this current study, we present a promising human RPE-porcine retina hybrid co-culture model that mimics some aspects of AMD and aids in understanding the relationship between RPE cells and PR. The first advantage of this model is the utilization of the avascular cone-rich zone in the porcine eye, which is more similar to the para-foveal cone-rich region of the human eye, than commonly used rodent models[6]. Secondly, this organotypic culture preserves the complex cell-cell interaction of the retina, including glia-neuron and RPE-retina interactions, which are important for maintaining retinal structure and function *ex-vivo*. Moreover, the porcine eyes that are more readily available as a by-product of the meat industry can be used for larger screening, and validation can be performed using human donor eyes, which are more rarely available. Indeed, we demonstrate that our model is also suitable for human RPE-human retinal explant co-culture.

The model is not intended to replicate AMD fully, but it is designed to answer specific questions concerning AMD pathology. Considering that the RPE cell component of the model can be genetically manipulated or exposed to AMD-related stressors prior to co-culture, the main purpose of the model is to be employed in understanding the impact that healthy/diseased RPE cells have on the neuroretina. The diseased RPE cells for this study were generated by dysregulation of FH, which is a known inhibitor of the complement system. One of the main genetic risks for AMD corresponds to the Y402H polymorphism in the FH protein, but the mechanism by which FH Y402H polymorphism contributes to AMD is not fully understood, due to the multi-functional properties of the FH protein [4]. When secreted, FH exerts its anti-complement activation role and the FH 402H variant impairs this activity on the surrounding extracellular matrix, leading to complement activation and inflammation, which are main features of AMD [4]. However, recently an additional intracellular role for FH, independent from complement regulation, has been shown in different cell types including RPE cells [17, 18]. In our previous studies [17, 21], we show that loss of intracellular FH in hTERT-RPE1 cells, *via CFH* silencing, affects several AMD-relevant features and pathways in RPE cells in a similar way as the *CFH* 402H variant has been described to affect iPS-RPE cells [19, 20]. FH loss in RPE cells, besides altering complement and inflammatory regulation, had a great impact on RPE cells homeostasis. In particular, *CFH* silenced RPE cells present decreased mitochondrial respiration and glycolysis, indicating a general reduction in metabolic capacity. Moreover, *CFH* silenced RPE cells show increased vulnerability to oxidative stress and increased oxidized lipids content, both features found in AMD. Therefore, the first goal of this study was to investigate whether FH deprivation in RPE cells is sufficient to have detrimental effects on the neuroretina, independently from systemic complement sources. Additionally, the aim was to identify the main cell types affected in the neuroretina and the main changes in the ONL, which is the first retinal layer affected in AMD.

First, we examined the effects that RPE deprived of FH have on the structure of the co-cultured retina. We observed a reduction in all three parameters – retinal thickness, ONL thickness and the number of rows in ONL, indicating that RPE cells deprived of FH are less able to support the co-cultured porcine retina, therefore inducing retinal degeneration. In a *CFH* deficient mouse model, a similar phenomenon was reported in terms of photoreceptor loss in aged mice and in this study, it has been postulated that this could be due to RPE dysfunction and retinal stress [37]. In other moue models of retinal degeneration, where RPE damage was induced by deletion of Superoxide Dismutase 2 (*Sod2*)[38], it was reported that the damage to the RPE cells precedes and induces photoreceptor loss. The reduction in thickness and the number of cell rows in the ONL in our model, clearly indicates that a loss of PR occurs. However, as for AMD, it is under debate whether rods or cones degenerate first. Within the PR, it is known that the rods and cones have diverse metabolism [39], which contributes to differential susceptibility to different stress conditions [40, 41]. In our study, we found that after 3 days in co-culture, FH deprivation in the RPE cells induced loss of only rod photoreceptors, since the number of cones was not significantly changed. A similar trend was observed in AMD patients, where rod degeneration precedes cone degeneration in the para-foveal regions in early stages of AMD [42]. This result implies that the rod dominant retinal regions are more susceptible to early degenerative changes in AMD patients and in this co-culture model. However, in some cases, the cones nuclei, which in physiological conditions, are aligned in one single row at the most outer layer, appear displaced. Displacement of cones nuclei towards the outer plexiform layer of the retina has been previously described in another model of retinal degeneration and in eyes from aged and AMD donors [43], where it has been suggested that displaced cones, presumably losing their synaptic contacts do not immediately undergo cell death; but lose their functionality.

We defined that RPE cells silenced for FH cause PR degeneration. As the next step, we aimed to identify the mechanism by which this damage occurs. Inflammation is considered a major player in retinal degeneration and considering that FH is part of the innate immune system as a complement regulator and that FH-deprived cells modify the microenvironment towards an inflammatory milieu [21], we investigated the role of inflammation at different levels. One of the major contributors for neuro-inflammation is the retinal microglial cells [44]. Upon excessive activation of microglial cells, the retina undergoes chronic inflammation, leading to retinal degeneration [45]. Our results showed that the loss of FH in the RPE cells did not impact the number and activation of the microglial cells. This result suggests a glial cell-independent inflammatory response in the porcine retina co-cultured with FH deprived RPE cells, as reported in other neuro-degenerative diseases [46]. Previous studies on retinal degeneration have suggested that photoreceptor loss can also be dependent on modification of Müller cell structure and metabolism [47]. However, no significant differences were observed in Müller cell activation, and this implies absence of reactive gliosis in this model of FH loss induced retinal degeneration. Most importantly, we show that exogenous application of FH in the co-culture media does not rescue the neuroretina and that additional C3 does not exacerbate the effects of FH loss or cause any damage in control cultures. These findings support the hypothesis that the neurodegeneration is dependent on the damage that loss of intracellular FH causes to the RPE cells.

RPE cells are responsible for the maintenance of retinal homeostasis, not only from a structural perspective, but also from a metabolic side[11]. Being the cell type between the neuroretina and the blood supply of the choroid, the RPE cells are responsible for nutrients transfer to the retina, and any impairment in RPE metabolism would affect the amount or type of nutrients that reach the retina, leading to alteration of retinal metabolism[16]. Moreover, the retina consists of a highly oxidative natural environment, which provides a hostile living condition for the PR and the RPE cells help protect the retina from oxidative stress. In the context of AMD, several studies reported impairment in metabolic homeostasis and oxidative balance and alterations in mitochondria metabolism and oxidative stress tolerance were also reported in the RPE cells deprived of FH used in this study. Therefore, we investigated the direct impact of those cells on the mitochondria of the cells present in the ONL, the retinal layer in direct contact with RPE cells. Interestingly, we observed a reduction in the mitochondrial protein cytochrome c, specifically in its reduced form. This kind of signature has been previously described in cardiomyocytes after treatment with H_2_O_2_, describing a situation of oxidative stress [31]. One of the consequences of oxidative stress in the retina is lipid peroxidation [48]. Of note lipid peroxidation products are one of the main components of drusen and FH dysregulation and FH 402H variant has already been associated with lipid peroxidation. Phospholipids are the main component of cellular membranes and DHA is the most abundant polyunsaturated fatty acid (PUFA) of the retina that fulfills critical functions in providing a conducive membrane environment for rhodopsin-based phototransduction with single-photon sensitivity, stabilization of photoreceptor outer segment membrane rim curvature, and fast synaptic vesicle fusion. DHA is also a main fatty acid chain of brain cardiolipin [49], an essential phospholipid constituent of the inner mitochondrial membranes (more than one-third of the acyl chains carry at least one DHA chain) and because of its wedge-like shape, cardiolipin stabilizes mitochondrial cristae shape [50, 51]. Therefore, oxidation of DHA has detrimental consequences on the neuroretina. When analyzing the lipid content *via* Raman in retinae cultured with FH-deprived cells, we observed indeed an increase in the oxidized fraction of DHA esters in the outer segments and inner segments of the ONL (PR), which could indicate a mitochondria and outer segments damage.

In summary, we developed a human RPE-porcine retina hybrid co-culture model of PR degeneration induced by a lack of FH in the RPE cells. Moreover, we provide evidence that the mechanism for this degeneration is likely dependent on intracellular FH in RPE cells and it is not directly mediated by inflammation. For instance, our data show that the retina, particularly the ONL, is affected at the level of mitochondria stability and oxidative stress balance. Thus, we developed a valuable and reliable preclinical model to study the pathogenesis of AMD, where the RPE component can be modified depending on the scientific question. For example, this model will be used in future studies to investigate the impact of iPSC-RPE carrying AMD high-risk genetic polymorphisms on the retina. Finally, our model can be used to screen potential therapeutic drugs to target AMD at the early stages of disease progression.

## Funding

Angela Armento is supported by the fortüne-Programm (project number 2640-0-0). This work was supported by donations from Jutta Emilie Paula Henny Granier and the Kerstan Foundation to Marius Ueffing and the Helmut Ecker Foundation to Simon J Clark.

## Acknowledgments

We are thankful to Karsten Schmidt (Retrotope, USA) for the advice on lipid metabolism.

## Conflicts of Interest

The authors declare no conflict of interest

## Notes

### Competing Interest Statement

The authors have declared no competing interest.

