## supplementary material for "FH loss in RPE cells causes retinal degeneration in a human RPE-porcine retinal explant co-culture model"

*Article*

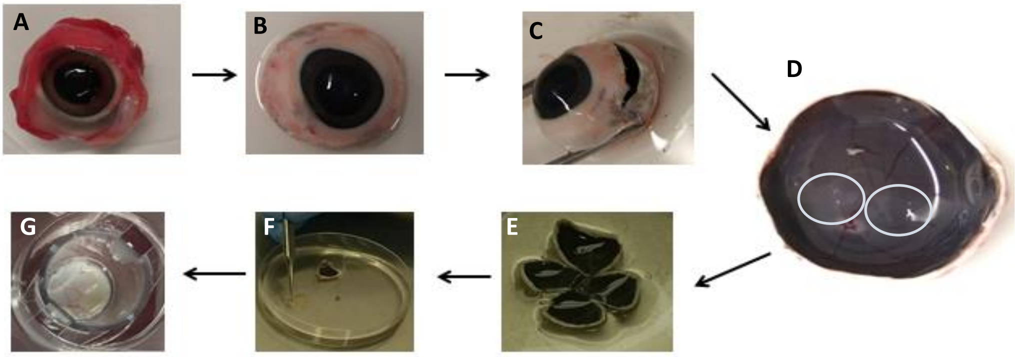


**Figure S1:** Preparation of porcine retinal explants for co-culture with RPE cells. **A.** Porcine eyecups were obtained from the slaughter house. **B.** Porcine eyecups after muscles and optic nerve removal. **C.** A small incision at about 5mm from the iris. **D.** The anterior segment of the eyecup including cornea, iris, lens and ciliary body were removed. The avascular cone-rich region of the retina was identified as indicated by white circles. **E.** Flattened eyecup. **F.** Retinal fragments gently peeled off from the RPE-choroid-sclera layers. **G.** Retinal explant placed on the transwell containing the RPE cells for co-culture.

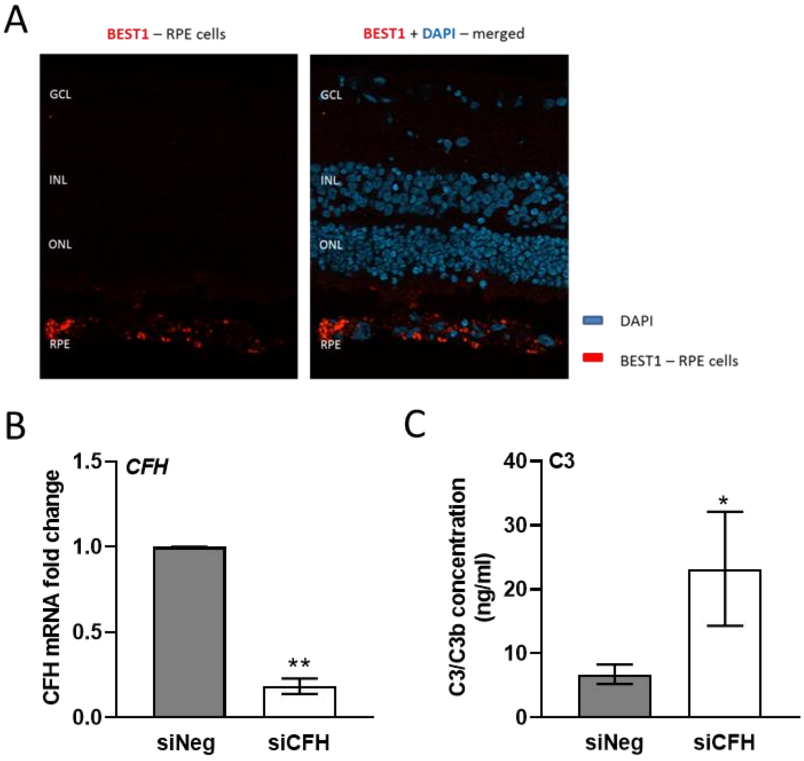


**Figure S2:** Characterization of hTERT-RPE1 cells used in the co-culture model. **A** porcine retinal explant in co-culture with hTERT-RPE1 cells. Cryosections were stained for RPE marker Best1 (red) and DAPI staining for nuclei. **B-C** Cell pellets and cell culture supernatants from Htert-RPE1 cells siNeg and siCFH were collected after 72 hours. **B** Evaluation of *CFH* expression after 72 hours by qRT-PCR analyses. Data are normalized to the housekeeping gene PRLP0 using Δ ΔCt methods. SEM is shown, n=4. **C** C3/C3b ELISA analyses of cell culture supernatants of hTERT-RPE1 cells. SEM is shown, n=4.

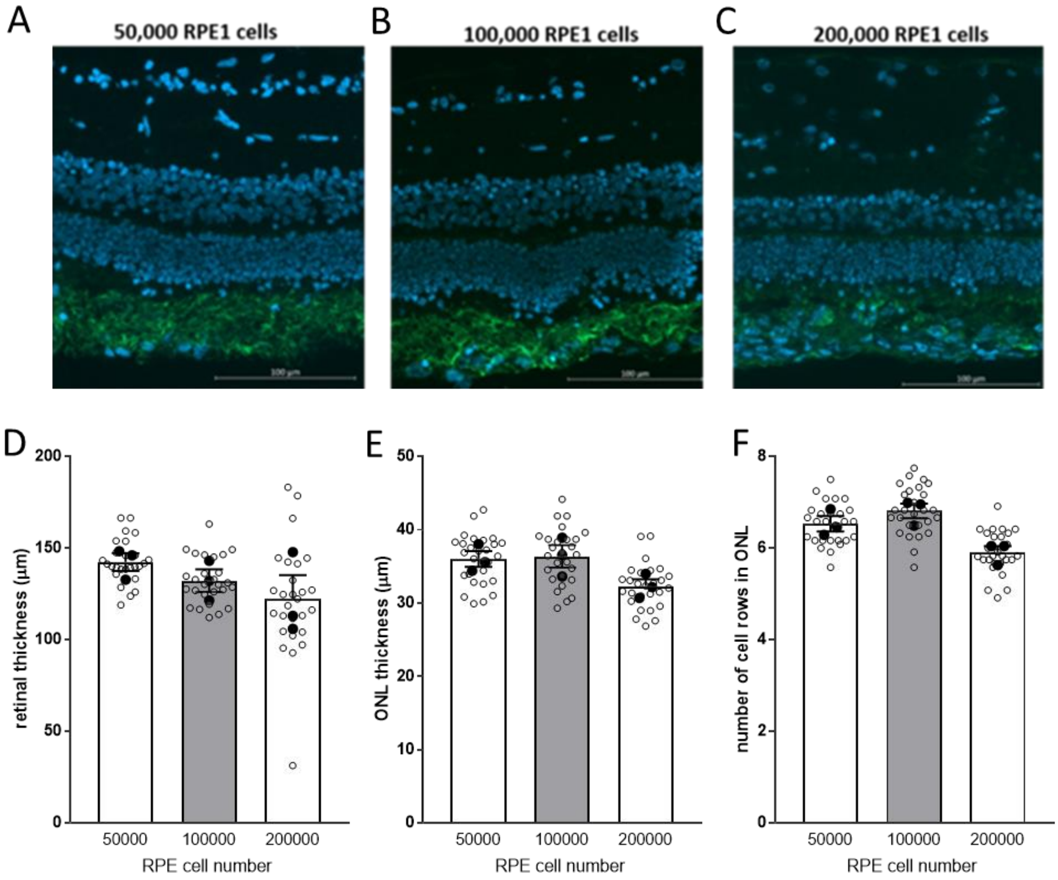


**Figure S3:** Effects of RPE cells seeding density on the co-cultured porcine retinae. Retinal explants were exposed to hTERT-RPE1 cells seeded at increasing cell density: **A** 50.000 cells, **B** 100.000 cells and **C** 200.000 cells. Retinal architecture parameters were assessed with the help of DAPI staining for nuclei. Quantification of the parameters are shown in D-F: retinal thickness (D), ONL thicknes (E), number of cell rows in the ONL (F). Mean ± SEM is shown. N=3.

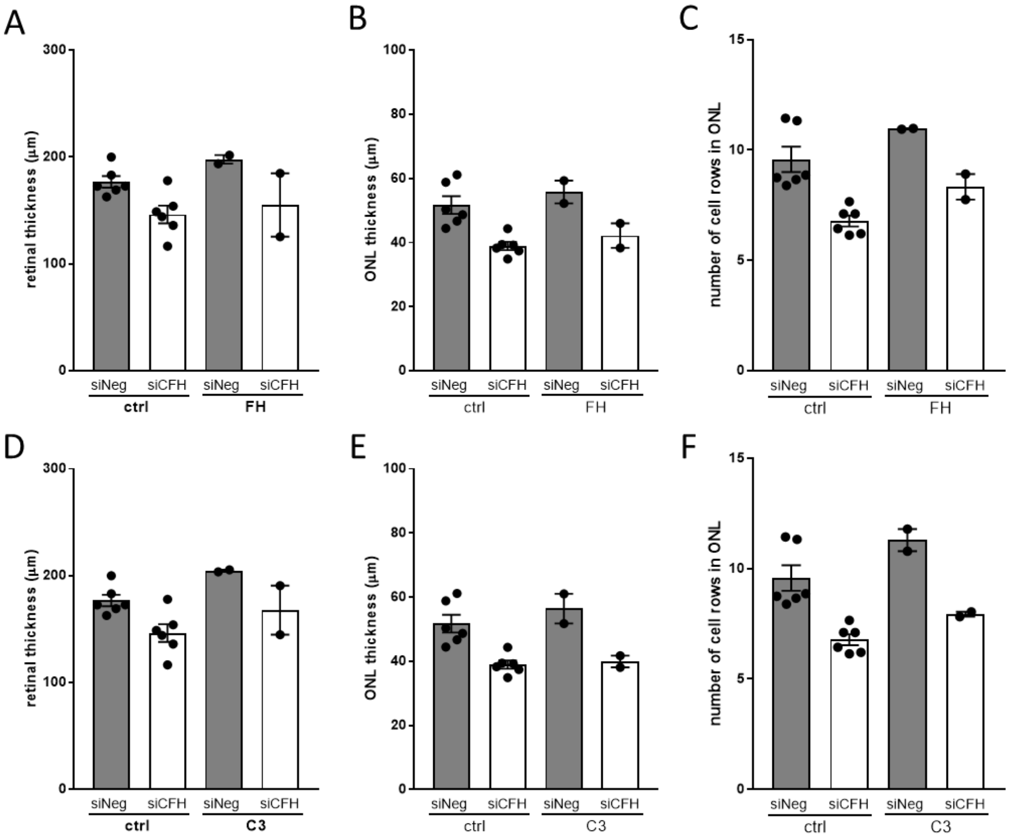


**Figure S4:** Exogenous FH or C3 does not rescue or exacerbate the effects of FH loss in RPE cells on the co-cultured porcine retinae. Retinal explants from the visual streak of porcine donor eyes are exposed to either siNeg (A) or siCFH (B) hTERT-RPE1 cells for 72 hours and medium was supplemented with purified FH or C3. Explants cryosections were stained with DAPI for nuclei and retinal architecture parameters were quantified: **A, D** retinal thickness, **B-E** ONL thickness and **C,F** number of cells in ONL. Abbreviations: GCL – Ganglion cell layer, INL – Inner nuclear layer, ONL – Outer nuclear layer, RPE – Retinal pigmented epithelium cells, siNeg – silencing negative control, si*CFH* – silencing *CFH*. Mean ± SEM is shown. N= 3 biological replicates. *$p \leq 0.05$; **$p \leq 0.005$.

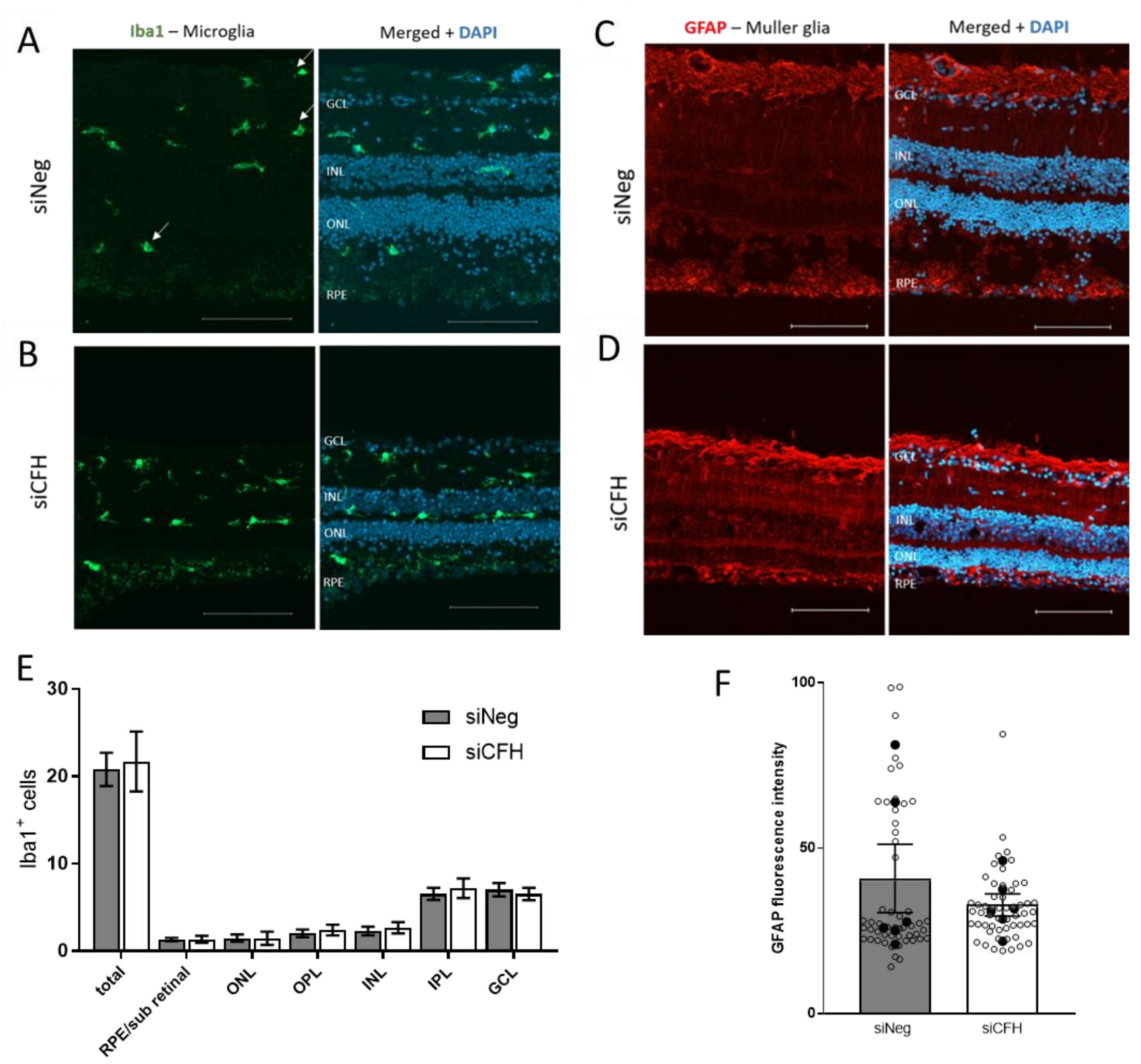


**Figure S5:** FH loss in RPE cells does not affect inflammatory cells on the co-cultured porcine retinae. Retinal explants from the visual streak of porcine donor eyes are exposed to either siNeg (A) or siCFH (B) hTERT-RPE1 cells for 72 hours. **A-B** Explants cryosections were stained for microglia marker Iba1 (green) and counterstaining was performed with DAPI for nuclei. **C-D** Explants cryosections were stained for Müller cells marker GFAP (red) and counterstaining was performed with DAPI for nuclei. **E** Quantification of Iba1 positive cells extrapolated from A-B. **F** quantification of GFAP fluorescence intensity extrapolated from C-D. Abbreviations: GCL – Ganglion cell layer, INL – Inner nuclear layer, ONL – Outer nuclear layer, RPE – Retinal pigmented epithelium cells, siNeg – silencing negative control, si*CFH* – silencing *CFH*. Mean ± SEM is shown. N= 6 biological replicates. *$p \le 0.05$; **$p \le 0.005$

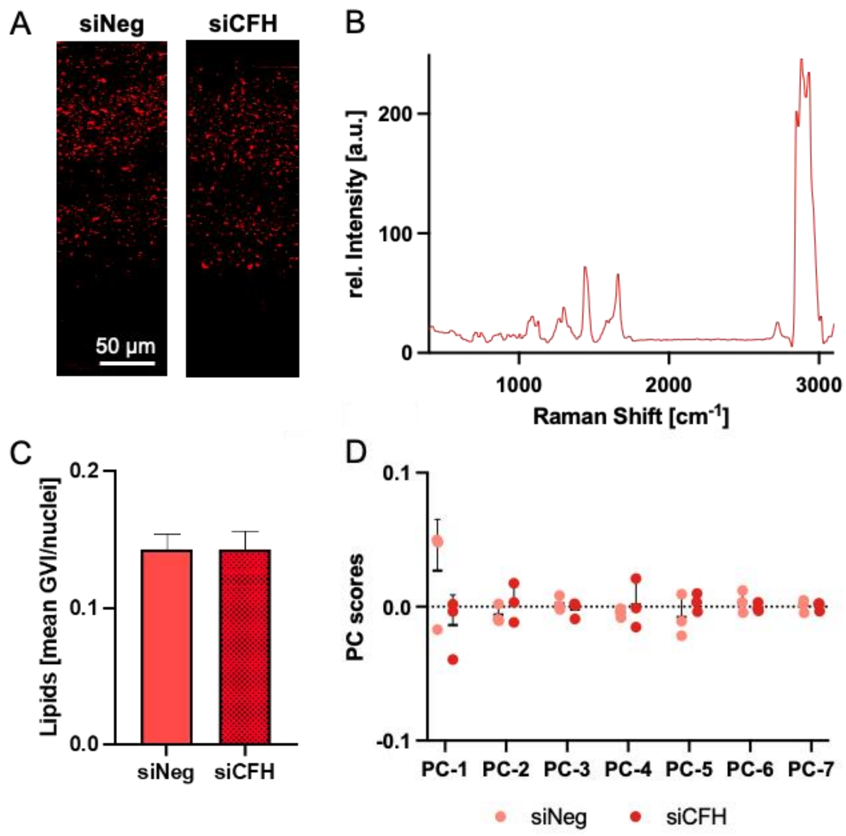


**Figure S6:** siCFH RPEs do not have an impact on the lipid expression and composition in the inner retina. **A-C** TCA identified a second lipid component localized in the inner retina (A) with a spectral signature assigned to phospholipids (B). Quantitative analysis by mean GVI/cell (C) did not show differences among both groups. **D** PCA of extracted lipid spectra expressed no significant difference in the lipid composition of the siNeg and siCFH group for any of the analyzed PCs. Data represent mean ± SD, n=3, scale bar – 50 µm
